# Chaperonin activity of *Plasmodium* prefoldin complex is essential to guard proteotoxic stress response and presents a new target for drug discovery

**DOI:** 10.1101/2022.09.17.508354

**Authors:** Rumaisha Shoaib, Vikash Kumar, Swati Garg, Monika Saini, Jyoti Kumari, Preeti Maurya, Aashima Gupta, Nutan Gupta, Harshita Singh, Pritee Verma, Ravi Jain, Shreeja Biswas, Ankita Behl, Mohammad Abid, Shailja Singh

**Affiliations:** Special Centre for Molecular Medicine, Jawaharlal Nehru University, New Delhi, Delhi, 110067; India; Medicinal Chemistry Laboratory, Department of Biosciences, Faculty of Natural Sciences, Jamia Millia Islamia, New Delhi, Delhi, 110025; India; Department of Life Sciences, Shiv Nadar University, Delhi NCR, Uttar Pradesh, 201314 India

## Abstract

The intraerythrocytic growth of malaria parasite is challenged by the presence of proteotoxic stress and intrinsically unstructured proteins in the cytoplasm due to formation of toxic heme during haemoglobin digestion. To overcome the unavoidable stress and maintain the cellular protein homeostasis, parasite encodes for a number of chaperones and co-chaperones. Here, we functionally characterize the *Plasmodium falciparum* prefoldins (*Pf*PFD1-6), a hexameric co-chaperone complex, for their role in protein homeostasis. We demonstrate that *Pf*PFD1-6 localise to cytosol of the parasite and the subunits perform an orchestrated interaction (-PFD3-PFD2-PFD1-PFD5-PFD6-PFD4-) to form an active jelly-fish like complex. Biperiden, an N-propylpiperidine analogue identified by chemotype search from FDA, strongly binds and restricts the formation of prefoldin complex and inhibited its interaction with the substrates, *Pf*MSP-1 and α-tubulin-I. Biperiden treatment potently inhibited the *in vitro* (IC_50_: 1μM) and *in vivo* growth of malaria parasite. Thus, this study provides novel virtues towards understanding the role of *Pf*PFDs in regulating protein homeostasis and opens new avenues for drug discovery against malaria.

## INTRODUCTION

The correct folding of proteins helps in maintaining the intracellular protein homeostasis which is vital for the conventional functioning of the cell (Tahmaz et al., 2022). Cell has a functional armamentarium of chaperones and co-chaperones that are involved in the surveillance of protein quality and conformation and assist in the correct folding of misfolded peptides (Hartl and Hayer-Hartl, 2002; Nielsen et al., 2014). The substrate recognition of the target mis-folded peptides is mediated by the co-chaperones which handover them to the respective chaperones (Millán-Zambrano and Chávez, 2014; Zhang et al., 2020).

Prefoldin (PFD) co-chaperone plays an essential role in maintaining cellular homeostasis under physiological and pathological conditions (Tahmaz et al., 2022). They are a heterohexameric molecular co-chaperone, which capture nascent polypeptide (e.g. actin and tubulin) and subsequently deliver it to TRiC/CCT complex for correct folding Prefoldin complex is present in archaea and eukaryotes and composed of six distinct subunits (PFD1-6) that form a jellyfish-like structure (Gestaut et al., 2019; Siegert et al., 2000; Vainberg et al., 1998). In archaea, PFD complex is comprised of two α and four β subunits, while in eukaryotes two α like (PFD3 and PFD5) and four β like (PFD1, PFD2, PFD4 and PFD6) subunits exist to form a complex. The formation of PFD complex protects it from ubiquitin-proteasome-mediated degradation (Miyazawa et al., 2011).

Chaperone activity of PFD plays a significant role in restricting the protein misfolding and thus reduced toxic aggregate formation in neurodegenerative diseases. Dysfunction of PFD in humans leads to the aggregation of proteins that trigger neurodegenerative diseases like Alzheimer’s (AD), Parkinson’s (PD), and Huntington’s diseases (HD) (Sörgjerd et al., 2013; Takano et al., 2014; Tashiro et al., 2013). Reports have shown that prefoldin not only inhibits α-synuclein aggregate formation but also assists autophagy-dependent degradation of α-synuclein aggregates (Takano et al., 2014).

PFD subunits especially promotes the actin and tubulin polymerization in the cytoplasm. For eg., in *Caenorhabditis elegans*, RNAi mediated knockdown of PFD leads to reduced α-tubulin levels; subsequently, microtubule polymerization rate is decreased (Locascio et al., 2013). Another report suggested that PFD-1 deficient mice have defects in the development and function of lymphocytes including abnormalities in cytoskeletal functions such as neuronal loss, and ciliary dyskinesia (Cao, 2016).

Recently, in *Plasmodium falciparum*, the accumulation of unfolded protein response in the cytoplasm of the parasite has been correlated with the artemisinin resistance (Mok et al., 2015). Furthermore, the proteome of *P. falciparum* is enriched in intrinsically unstructured proteins and since, the structural integrity of proteins is important for the survival of cell, the 2% of the parasite’s highly reduced genome is encoding for the chaperones (Feng et al., 2006). Thus, suggesting the importance of molecular chaperones for the parasite in guarding the correct protein folding.

The fundamental research on PFD subunits have demonstrated their enormous potential and importance in maintaining cellular proteastasis and survival. However, the complete lack of understanding of PFD complex in malaria parasite remains a big void in the understanding of functional chaperones and co-chaperones. This foundational study on *Plasmodium falciparum* PFD (*Pf*PFD) subunits completely characterizes the functional and biochemical roles of this complex. We demonstrate that the six subunits of *Pf*PFD are expressed as a complex in the cytoplasm of the trophozoite and schizont stages of *P. falciparum*. Further, molecular interactions within the complex were investigated using microscale thermophoresis and demonstrate that the orchestrated organization of *Pf*PFD subunits forms the active PFD complex. Additionally, we demonstrate that *Pf*PFD complex interacts with a-tubulin through *Pf*PFD2 both *in vivo* and *in vitro* indicating that *Pf*α-tubulin-I is one of the substrates of *Pf*PFD complex. Interrogation of chemotypes from FDA against *Pf*PFD, identified Biperiden (BPD) as a potential inhibitor of the complex. BPD restricts the formation of *Pf*PFD complex *in vitro* and demonstrate potent anti-malarial activity. Treatment with BPD disrupted the PFD complex and induced degradation of its substrate tubulin. Owing to the fact that PFD interacts with unstructured proteins and merozoite surface proteins are highly rich in intrinsically unstructured regions, we demonstrate that inhibition of the *Pf*PFD complex by BPD treatment leads to degradation of *Pf*MSP1 *in vivo*. This study describes that the fundamental functions of *Pf*PFDs are conserved with other PFDs however, the *Pf*PFDs have diverged and specialized themselves to interact with the unique substrates present in the P. falciparum proteome. Thus *Pf*PFDs provide a unique opportunity for drug development against malaria parasite.

## RESULTS

### Orchestrated assembly of the *Plasmodium falciparum* Prefoldin hetero-hexamer

*Plasmodium falciparum* 3D7*(Pf*3D7) PFD subunits, namely *Pf*PFD1 (20 kDa), *Pf*PFD2 (19 kDa), *Pf*PFD3 (24 kDa), *Pf*PFD4 (17 kDa), *Pf*PFD5 (30 kDa) and *Pf*PFD6 (16 kDa), were cloned in pET-28a (+) and over-expressed in *E. coli* strain BL21-DE3. Recombinant 6x-His-*Pf*PFD (r*Pf*PFD) subunits were purified using Ni-NTA resin, as described previously (Kumar et al., 2022a) (Fig. S1). Polyclonal antisera were raised against all *Pf*PFDs in male BALB/c mice. The endpoint titers and antibody specificity of raised antisera were determined before using them for future experiments.

Individual structural models of *Pf*PFD subunits 1-6 were generated using I-TASSER (Zhang, 2008). The top threading templates used by I-TASSER are as follows: Prefoldin from *Pyrococcus horikoshii* OT3, chain C (PDB ID: 2ZDI; for *Pf*PFD1, *Pf*PFD3, and *Pf*PFD5); and, Prefoldin beta subunit from *Thermococcus* strain KS-1, chain A (PDB ID: 2ZQM; for *Pf*PFD2, *Pf*PFD4 and *Pf*PFD6) (Kida et al., 2008; Ohtaki et al., 2008), and set to submit to generate *Pf*PFD hetero-hexameric complex structure by using X-Ray diffraction-based structural model of human TRiC-PFD complex as a suitable template (Gestaut et al., 2019). After optimal rigid-body superimposition of the generated structural model of *Pf*PFD complex with *Hs*PFD, the overall Root-Mean-Square Deviation (RMSD) value of the C-alpha atomic co-ordinates was found to be 1.45 Å, suggesting a reliable 3D structure. Assessment of the stereochemical quality and accuracy of the generated structural model of *Pf*PFD complex displayed 91.8% of amino acid residues lying in the most favoured (core) regions, with 6.8%, 0.9%, and 0.5% residues in additional allowed, generously allowed, and disallowed regions of Ramachandran plot, respectively.

*Pf*PFD structural model revealed a strong resemblance with its counterparts from other eukaryotes, forming a hetero-hexamer with a jellyfish-like architecture composed of two canonical classes of subunits: α (PFD subunits: 3 and 5) and β (subunits: 1, 2, 4, and 6), arranged in the following manner -PFD3-PFD2-PFD1-PFD5-PFD6-PFD4-(Gestaut et al., 2019; Siegert et al., 2000) (Fig. 1A). The core body of the *Pf*PFD complex was found to consist of a double beta-barrel assembly, with six long tentacle-like coiled coils protruding from it in a regularly structured fashion. The comparable RMSD value and Ramachandran plot characteristics confirmed the reliability of the *Pf*PFD hetero-hexameric complex to be taken further for *in silico* and *in vitro* interaction analysis, and inhibition studies.

**Figure 1:**
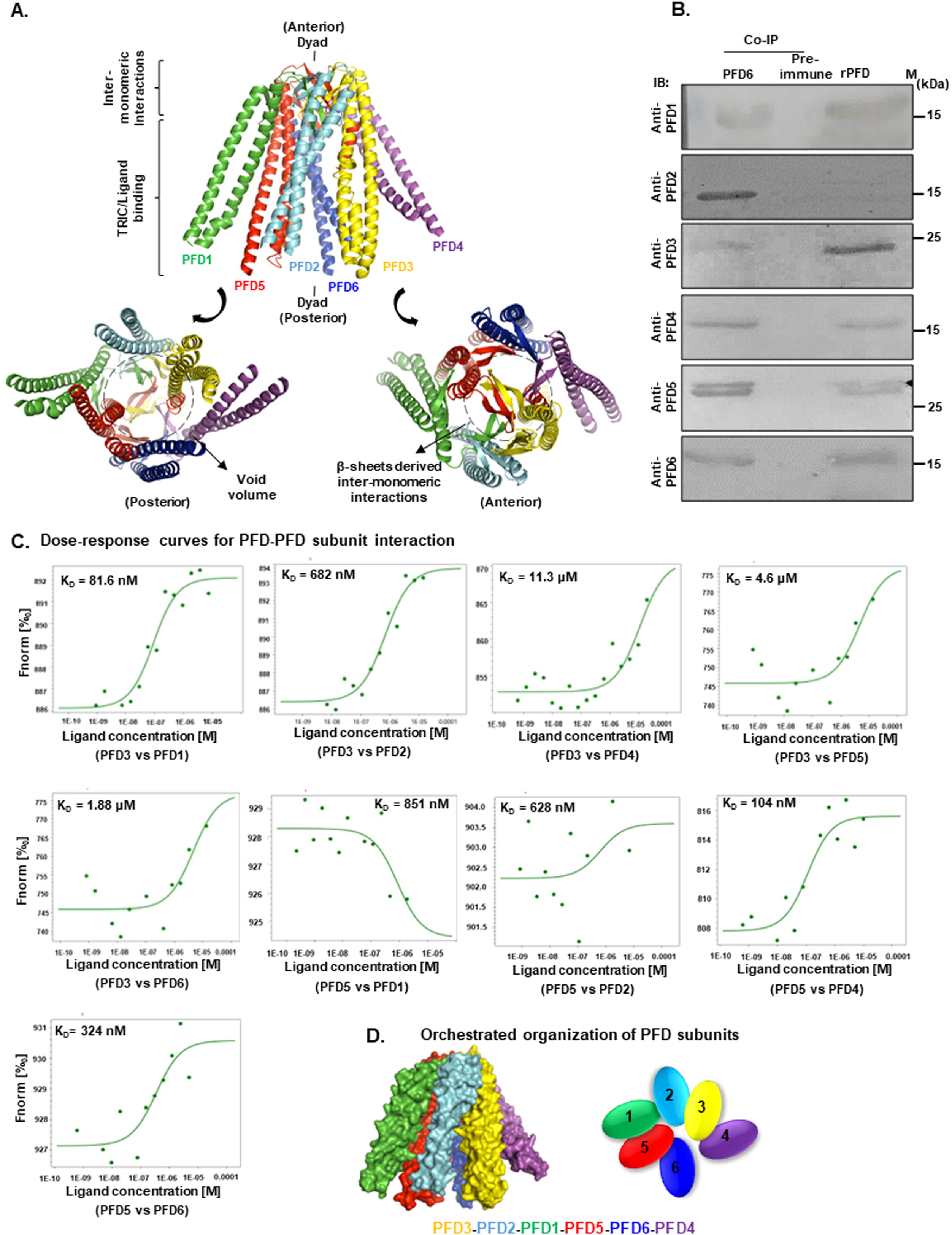
Assembly of hetero-hexameric PfPFD complex in vitro and in vivo. **(A) Overall architecture of the *Pf*PFD-hexameric complex**. Structural models of *Pf*PFD subunits 1-6 were generated using I-TASSER, and *Pf*PFD hetero-hexameric complex structure was generated using human TRiC-PFD complex (PDB ID: 6NR8) as a template. Upon rigid-body superimposition of both the complexes, the overall RMSD value of the C-alpha atomic co-ordinates was found to be 1.45 Å, suggesting a reliable 3D structure. **(B)Existence of prefoldin subunits as a complex confirmed by co-immunoprecipitation**. Anti-*Pf*PFD6 antisera was cross-linked to amino coupling plus resin followed by incubation with parasite lysate. Beads bound fractions were eluted, resolved on 12 % SDS PAGE and subjected to Western blot analysis using anti*-Pf*PFD1, anti-*Pf*PFD2, anti-*Pf*PFD3, anti-*Pf*PFD4, anti-*Pf*PFD5, and anti-*Pf*PFD6 antibodies. **(C) One-to-one interaction study of prefoldin subunits by Microscale Thermophoresis**. 10 μM purified recombinant *Pf*PFD3 and *Pf*PFD5 were labelled with protein labelling dye RED-NHS (L001, NanoTemper technologies, Germany). Varying concentrations of unlabelled proteins i.e., *Pf*PFD1 and *Pf*PFD2, *Pf*PFD4, *Pf*PFD5, *Pf*PFD6 for labelled *Pf*PFD3 and, *Pf*PFD1 and *Pf*PFD2, *Pf*PFD3, *Pf*PFD4 and *Pf*PFD6 for labelled *Pf*PFD5 were titrated in 1:1 dilution. All measurements were carried out in samples filled capillaries. Sigmoidal curve generated K_D_ of 81.6 nM, 682 nM, 11.3 μM, 4.6 μM, 1.88 μM, 851 nM, 628 nM, 104 nM, 324 nM for *Pf*PFD1-*Pf*PFD3, *Pf*PFD2-*Pf*PFD3, *Pf*PFD4- *Pf*PFD3, *Pf*PFD5-*Pf*PFD3, *Pf*PFD6-*Pf*PFD3, *Pf*PFD1-*Pf*PFD5, *Pf*PFD2-*Pf*PFD5, *Pf*PFD4-*Pf*PFD5, *Pf*PFD6-*Pf*PFD5 pairs, respectively.

Protein-protein interactions (PPI) are crucial in determining the target protein function and drugability of molecules. In this regard, binding affinities among the r*Pf*PFD subunits were evaluated by MST analyses, using Monolith NT.115 instrument (NanoTemper Technologies, Munich, Germany). MST relies on binding-induced changes in thermophoretic mobility, which depends on several molecular properties including particle charge, size, conformation, hydration state, and solvation entropy. Thus, under constant buffer conditions, the thermophoresis of unbound proteins typically differs from the thermophoresis of proteins bound to their interaction partners. The thermophoretic movement of a fluorescently labeled protein is measured by monitoring the fluorescence distribution.

In eukaryotes and archaea, PFD2 and PFD6 interact with PFD3 and PFD5, respectively to generate sub-complexes. PFD1 and PFD4 are recruited and assembled by these sub-complexes to form a complex (Liang et al., 2020). Based on the earlier reference, we labeled *Pf*PFD3 and *Pf*PFD5 and demonstrated their interaction with other PFD subunits. The concentration of the labeled protein was kept constant at 20 nM and titrated with the interacting subunits. The binding affinity (K_D_) was revealed by steady-state analysis, which was obtained by plotting response at equilibrium as a function of unlabelled protein concentration (*Pf*PFD1: 7.5 μM, *Pf*PFD2: 28 μM, *Pf*PFD3: 20 μM, *Pf*PFD4: 20 μM, *Pf*PFD5: 13.4 μM, and *Pf*PFD: 6 20 μM). K_D_ values were determined to be 81.6 nM (*Pf*PFD3-*Pf*PFD1), 682 nM (*Pf*PFD3-*Pf*PFD2), 11.3 μM (*Pf*PFD3-*Pf*PFD4), 4.6 μM (*Pf*PFD3-*Pf*PFD5), and 1.88 μM (*Pf*PFD3-*Pf*PFD6), indicating good binding strength between *Pf*PFD3-*Pf*PFD1 and *Pf*PFD3-*Pf*PFD2, in comparison to *PfPFD*3-*Pf*PFD4, *Pf*PFD3-*Pf*PFD5 and *Pf*PFD3-*Pf*PFD6 (Fig. 1 B). Similarly, K_D_ values were determined to be 851 nM (*Pf*PFD5-*Pf*PFD1), 628 nM (*Pf*PFD5-*Pf*PFD2), 104 μM (*Pf*PFD5-*Pf*PFD4), and 324 nM (*Pf*PFD5-*Pf*PFD6), indicating higher binding affinity between *Pf*PFD5-*Pf*PFD4 and *Pf*PFD5-*Pf*PFD6, as opposed to *Pf*PFD5-*Pf*PFD1 and *Pf*PFD5-*Pf*PFD2 (Fig 1 C).

*Pf*PFD complexation was confirmed with co-immunoprecipitation assay (co-IP), in which *Pf*PFD6 antiserum was cross-linked to AminoLink plus Coupling Resin followed by incubation with the parasite lysate. Eluted fractions were resolved on SDS-PAGE, followed by western blotting with the respective *Pf*PFD antisera. Each blot showed a respective band of the *Pf*PFD subunits (Fig 1 B). As a negative control, pre-immune serum was cross-linked to the resin, while r*Pf*PFD subunits were used as a positive control. These findings suggested that *Pf*PFD subunits are present in a complex form as their counterparts in other species.

### Expression and localization of PfPFD subunits in the blood stage of malaria parasite

Semi-quantitative RT-PCR revealed that the transcripts encoding *Pf*PFD subunits are expressed throughout the intra-erythrocytic life cycle of the parasite. Expression of *Pf*PFD1 and *Pf*PFD6 was observed to be higher at the trophozoite and schizont stages compared to the ring stage, whereas *Pf*PFD4 expression was found to be higher at the trophozoite stage (Fig 2A). As a positive control, transcript encoding for 18s rRNA was amplified using *pf18s* specific primers. Furthermore, using the respective in-house developed polyclonal antisera, western blot analysis of the native *Pf*PFD subunits in the parasite lysates derived from all three intra-erythrocytic stages corroborated the expression of *Pf*PFD subunits in the parasite. At trophozoite and schizont stages, distinct protein bands with anticipated molecular weights were identified for all *Pf*PFD subunits (18.4 kDa: *Pf*PFD1, 16.6 kDa: *Pf*PFD2, 22.6 kDa: *Pf*PFD3, 15.3 kDa: *Pf*PFD4, 29.1 kDa: *Pf*PFD5, and 13.6 kDa *Pf*PFD6) (Fig. 2C). At the ring stage, no protein was found. Tubulin was taken as a loading control. Furthermore, as negative controls, no protein was found in the *E. coli* lysate or uninfected RBCs (both insoluble and soluble fractions) (Fig. S2). A comparable blot probed with pre-immune serum indicated the absence of the protein (data not shown). These findings suggest that *Pf*PFD subunits are expressed at both the transcript and protein levels throughout the asexual intra-erythrocytic stages of the parasite.

**Figure 2:**
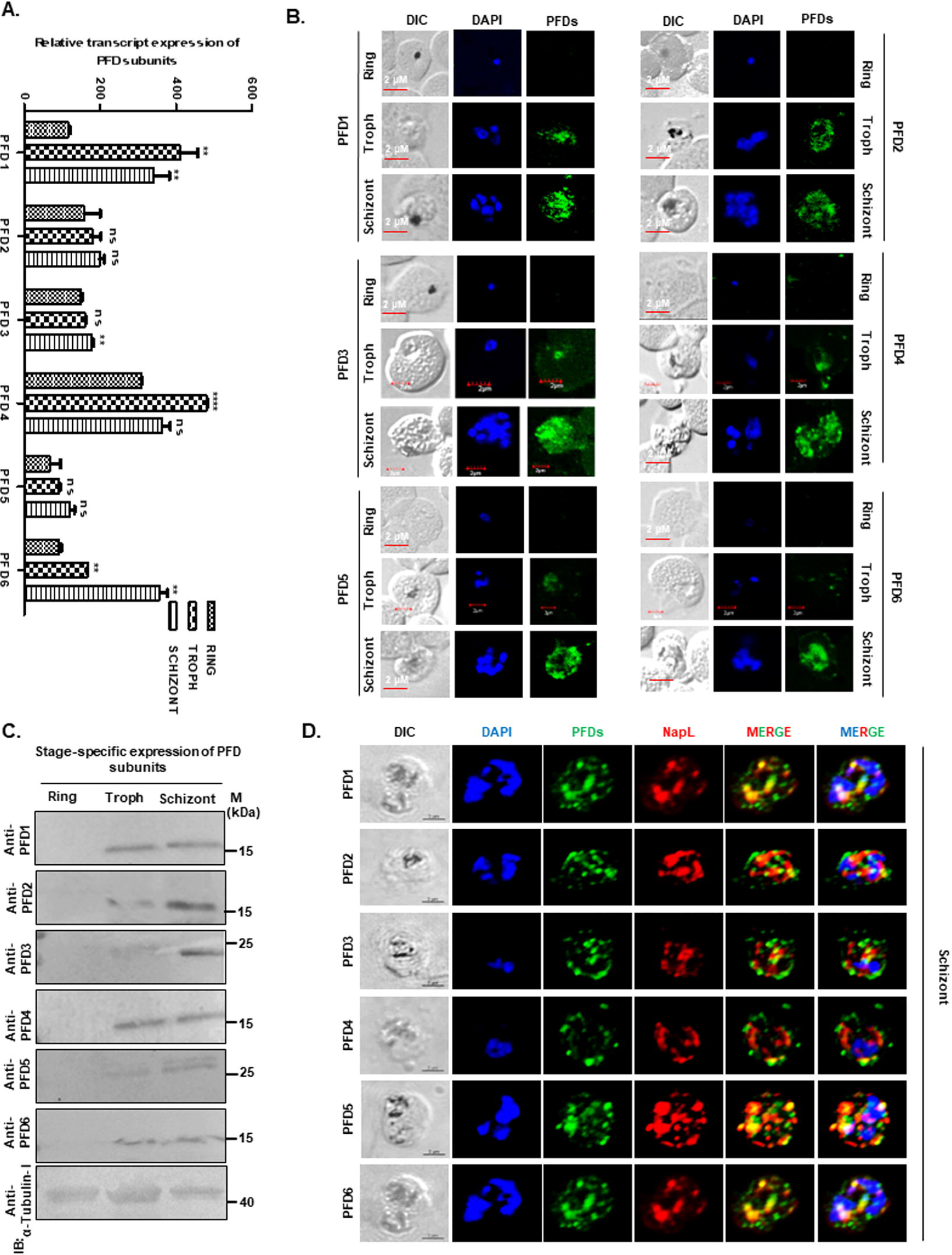
Expression and localization of PfPFDs in blood stage of *P. falciparum*. **(A) Stage-specific expression analyses of prefoldin subunits at transcript level**. Transcript levels of prefoldin subunits at three different asexual stages of *Pf*3D7 genome were measured by real-time PCR using specific primer sets. All values are presented as relative copy numbers. (^ns^*p*>0.05; ^*^*p*<0.05; ^**^*p*<0.01; ^***^*p*<0.001) **(B) (i) Localization analyses of prefoldin subunits at asexual stages of *Pf*3D7**. IFA images of *Pf*PFD1, *Pf*PFD2, *Pf*PFD3, *Pf*PFD4, *Pf*PFD5 and *Pf*PFD6 at three different asexual blood stages of *Pf*3D7. Methanol-fixed smears of infected erythrocytes were stained with anti-prefoldin subunits antisera (1:250) followed by incubation with Alexa Fluor conjugated secondary antibodies (1:200; Alexa Fluor 488, green color). DIC: differential interference contrast image, DAPI: nuclear staining using 4^’^,6-diamidino-2-phenylindole (blue); *Pf*PFD1, *Pf*PFD2, *Pf*PFD3, *Pf*PFD4, *Pf*PFD5, and *Pf*PFD6: mouse anti-*Pf*PFD1, anti-*Pf*PFD2, anti-*Pf*PFD3, anti-*Pf*PFD4, anti-*Pf*PFD5, and anti-*Pf*PFD6 antibodies, respectively (green) **(C) Stage-specific expression analyses of prefoldin subunits at protein level**. Blot showing the expression of *Pf*PFD1, *Pf*PFD2, *Pf*PFD3, *Pf*PFD4, *Pf*PFD5, and *Pf*PFD6 at different asexual stages of parasite development. **(D) Co-localization of prefoldin subunits with cytosolic marker NapL**. Expression and co-localization analyses of prefoldin subunits of *Pf*PFD1, *Pf*PFD2, *Pf*PFD3, *Pf*PFD4, *Pf*PFD5 and *Pf*PFD6 with *Pf*NapL at schizont stage of parasite life cycle. Methanol fixed infected RBC smears were stained with anti-prefoldin subunits antibodies (1:200) and anti-*Pf*NapL antibodies (1:250) followed by incubation with Alexa Fluor-conjugated secondary antibodies (Alexa Fluor 488, green; Alexa Fluor 546, red). DIC: differential interference contrast image, DAPI: nuclear staining using 40, 6-diamidino-2-phenylindole (blue); prefoldin subunits: mouse anti-prefoldin subunits antisera (green); *Pf*NapL: anti- *Pf*NapL antibody (red); merge: overlay of prefoldin subunits proteins with *Pf*NapL.

Immunofluorescence assays (IFAs) performed to visualize the subcellular expression and localization of the *Pf*PFD subunits during the asexual stages of the malaria parasite revealed that expression of all *Pf*PFD subunits begins at the trophozoite stage (Fig. 2B). The expression of *Pf*PFD subunits appeared to be confined to the parasite cytoplasm. *Pf*PFD subunits were co-localized with *Pf*NapL, a previously known cytosolic protein, to corroborate its cytosolic localization. At the schizont stage, there was significant overlapping of *Pf*PFD subunits (green) and *Pf*NapL (red), indicating their coexistence in the cytoplasm of the malaria parasite (Fig 2D).

### Functional interaction of *PfP*FD subunits with α-tubulin-I and PfMSP-1

Previous reports in eukaryotes indicate that PFD binds to cytoskeletal proteins (Abe et al., 2013; Povarova et al., 2014; Zhang et al., 2016). Protein-protein interaction data available in the PlasmoDB database also indicate the interaction of *Pf*PFD2 with *Pf*α-tubulin-I. To validate the interaction, preliminary screening was carried out using semi-quantitative ELISA, in which *Pf*PFD subunits 1-6 upon titration with *Pf*α-tubulin-I depicted the interaction in a concentration-dependent manner (Fig. 3A). To quantify the interaction strength, SPR-based interaction analysis was performed in which upon titrating the immobilized *Pf*PFD2 with *Pf*α-tubulin-I, K_D_ value was found to be 1.9◻×◻10^−7^ M (Fig. 3B). SPR analysis was also performed by titrating the immobilized *Pf*PFD1 and *Pf*PFD3 with *Pf*α-tubulin-I, which exhibited no interaction (Fig.S2 i,ii), demonstrating that *Pf*α-tubulin-I specifically binds to *Pf*PFD2, but not to *Pf*PFD1 or *Pf*PFD3. Further, to evaluate the expression and co-localization of *Pf*PFD2 with α-tubulin-I in the intra-erythrocytic stages of the parasite, an immunofluorescence assay was performed, which depicted that the two proteins co-localize at the trophozoite and schizont stages (Fig. 3C). The interaction of *Pf*PFD2 with α-tubulin-I was confirmed with co-IP, in which *Pf*PFD2 antiserum was cross-linked to the AminoLink plus Coupling Resin, followed by incubation with the parasite lysate prepared from the mix-stage parasite population. The desired protein band of α-tubulin-I was observed in the eluted fraction (Fig. 3D (i)). Similarly, reverse co-IP, in which *Pf*α-tubulin-I antisera was cross-linked to the resin, followed by incubation with the parasite lysate confirmed the interaction between the two proteins (Fig. 3D (ii)). Collectively, these findings point to a possible interaction between *Pf*PFD2 and α-tubulin-I. Similarly, co-IP indicated the interaction of *Pf*PFD2 and *Pf*PFD6 with PfMSP-1 and α-tubulin-I, respectively (Fig. 3D (iii) and (iv)).

**Figure 3.**
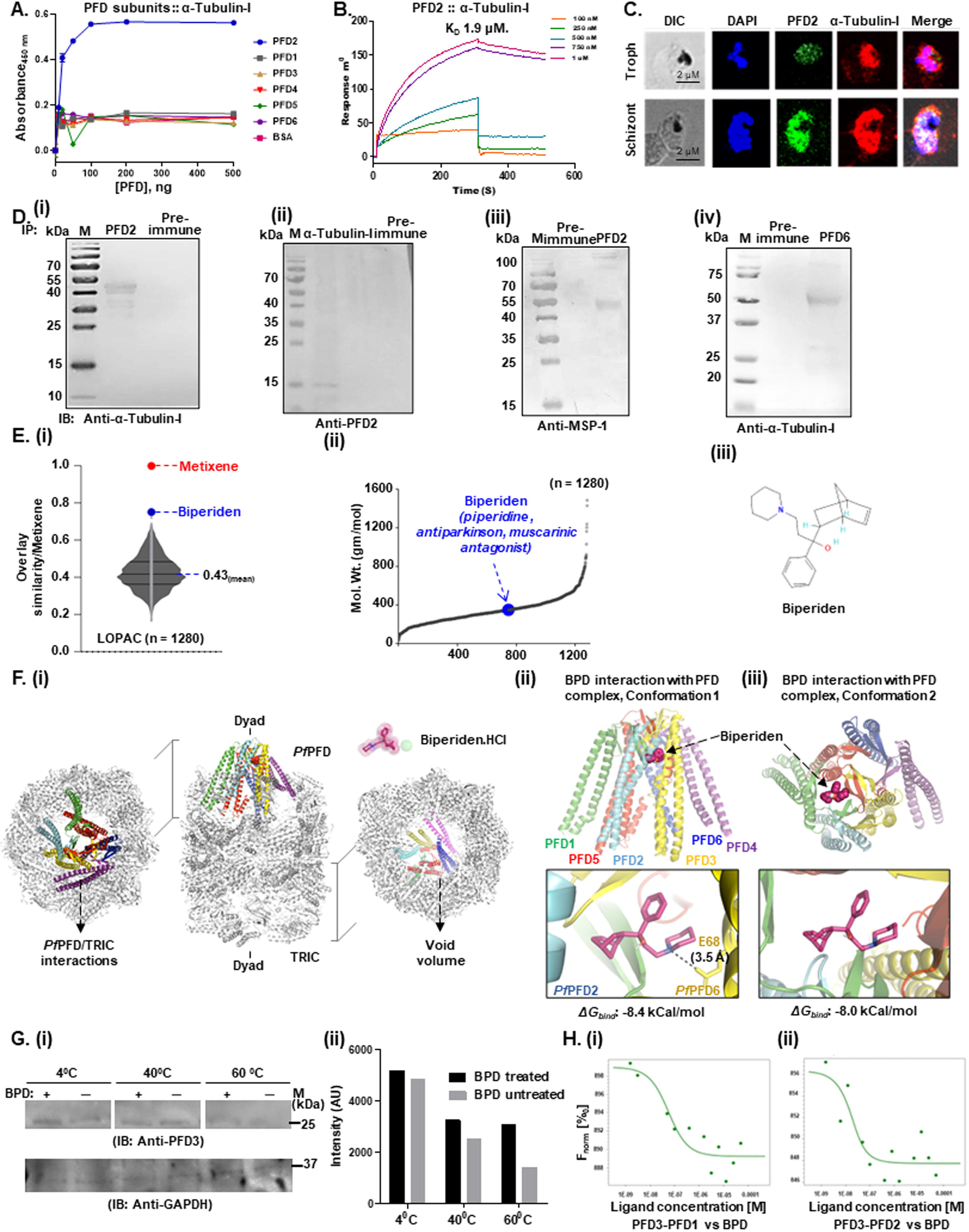
Functional interaction of PfPFD complex with its substrates and identification of Biperiden as its inhibitor. **(A) Monitoring the interaction of *Pf*PFD2 with α-tubulin-I by ELISA** Concentration dependent binding curves of *Pf*PFD1, *Pf*PFD2, *Pf*PFD3, *Pf*PFD4, *Pf*PFD5 and *Pf*PFD6 with α-tubulin-I where y-axis represents absorbance at 490 nm and x-axis denotes concentration of prefoldins. Error bar represent standard deviation among the three replicates. Orange line depicts the binding of *Pf*PFD2 with α-tubulin-I whereas blue, green, red, grey and yellow depicts binding of *Pf*PFD1, *Pf*PFD3, *Pf*PFD4, *Pf*PFD5 and *Pf*PFD6 with α-tubulin-I. **(B) Monitoring the interaction of *Pf*PFD-2 with α-tubulin-I by SPR**. Purified recombinant prefoldin subunits (25 μM) were immobilized on gold sensor chip followed by titration with varying concentrations of α-tubulin-I. Net dissociation constant graph generated K_D_ 1.9 μM. **(C) Co-localization of *Pf*PFD2 with α-tubulin-I**. Expression and co-localization analysis of *Pf*PFD2 with α-tubulin-I at trophozoite and schizont stages of parasite life cycle. Methanol fixed infected RBC smears were stained with anti-*Pf*PFD2 (1:200) and anti-α-tubulin-I antibodies (1:250) followed by incubation with Alexa Fluor-conjugated secondary antibodies (Alexa Fluor 488, green; Alexa Fluor 546, red). DIC: differential interference contrast image, DAPI: nuclear staining using 40, 6-diamidino-2-phenylindole (blue); *Pf*PFD2: mouse anti-*Pf*PFD2 sera (green); α-tubulin-I: anti-α-tubulin-I antibody (red); merge: overlay of *Pf*PFD2 proteins with α-tubulin-I. **(D) Interaction analyses of prefoldin subunits by Co-immunoprecipitation assays. (i)** Anti-*Pf*PFD2 sera was cross-linked to amino coupling plus resin followed by incubation with parasite lysate. Beads bound fractions were eluted, resolved on 12 % SDS-PAGE and subjected to Western blot analysis using anti-α-tubulin-I. **(ii)** Anti-α-tubulin-I sera were cross-linked to amino coupling plus resin followed by incubation with parasite lysate. Beads bound fractions were eluted, resolved on 12 % SDS-PAGE and subjected to Western blot analysis using anti-*Pf*PFD2 sera. **(iii)** Anti-*Pf*PFD6 sera was cross-linked to amino coupling plus resin followed by incubation with parasite lysate. Beads bound fractions were eluted, resolved on 12 % SDS-PAGE and subjected to Western blot analysis using anti-α-tubulin-I sera. **(iv)** Anti-*Pf*PFD2 antisera were cross-linked to amino coupling plus resin followed by incubation with parasite lysate. Beads bound fractions were eluted, resolved on 12 % SDS-PAGE and subjected to Western blot analysis using anti-MSP-I sera. **(E) Biperiden (BPD) as a possible inhibitor of *Pf*PFD protein-folding activity. (i)** LOPAC^®1280^ library was screened based on structural similarities with one of the potent protein folding activity of ribosomes (PFAR) inhibitors, Metixene. **(ii, iii)** BPD was identified with an overlay structural similarity index of 0.75. **(F) Possible architecture of the *Pf*PFD-BPD-TRiC (i)** The complex was generated using the modeled structure of the *Pf*PFD complex, and the human TRiC complex as a representative of *Pf*TRiC. BPD was found to interact with *Pf*PFD complex via two different conformations. **(ii)** In conformation 1, BPD engaged at the interface of *Pf*PFD subunits 2 and 6, with free binding energy (*ΔG_bind_*) of −8.4 kCal/mol. **(iii)** In conformation 2, BPD was found to interact with the double beta-barrel assembly of the *Pf*PFD complex, with *ΔG_bind_* of −8.0 kCal/mol. **(G) Identification of prefoldin as a target of BPD in parasite cell lysate**. The immune-based CETSA analysis of *Pf*PFD3 thermostability in the presence and absence of BPD. BPD treated, saponized parasite lysate was RIPA lysed and soluble protein fraction was pre-incubated with 5 μM BPD for 30 minutes, BPD untreated parasite was taken as control. **(i)** After thermal denaturation at 40 and 60°C, samples were processed for Western blotting and probed with anti-*Pf*PFD3 sera. As a reference point, the protein’s intensity at 4°C was used**(ii).**Using ImageJ software, the densitometry quantification of the relevant immunoblot was carried out and presented as a bar graph. Blotting with anti-GAPDH was used as a loading control. **(H) BPD competes with one prefoldin subunit for binding to the other**. Interactions of one prefoldin subunit with other prefoldin subunits were evaluated through MST. The K_D_ constant of 81.6 nM and 682 nM were obtained in case of interaction of *Pf*PFD3 with *Pf*PFD1 and *Pf*PFD2, respectively. An increasing MST signal (F_norm_ [%◻] starting at 886.1 units to 892.1 units for *Pf*PFD1, 886.4 to 893.8 for PFD2) was observed with increasing concentration of *Pf*PFD1 and *Pf*PFD2 yielding a sigmoidal curve (as in Figure 1C). **(ii)** Competition with ligand BPD as a competing molecule interfering with the interaction of *Pf*PFD3 with *Pf*PFD1 and *Pf*PFD2 displayed (F_norm_ [%◻] starting at **(i)** 893 units to 888.1 units for *Pf*PFD1**(ii)** and 894.5 to 887.6 for *Pf*PFD2) with K_D_ value 1.99 uM and 9.6 nM, respectively with increasing concentration of the ligand.

### Identification of Biperiden as the potential inhibitor of *Pf*PFD chaperonin activity

The *in silico* therapeutic repositioning approach which we adopted to screen the LOPAC^®1280^ library based on similarities with one of the potent PFAR inhibitors, Metixene, generated Biperiden, with an overlay structural similarity index of 0.75. Similar to Metixene, Biperiden (IUPAC name: 1-{bicyclo[2.2.1]hept-5-en-2-yl}-1-phenyl-3-(piperidin-1-yl)propan-1-ol) is a member of piperidines, and has a role as an antiparkinson drug and a muscarinic antagonist. In a recent investigation, Biperiden along with 16 other FDA-approved drugs, were identified to harbor anti-prion activity (Bamia et al., 2021). In the study, seven out of the 17 compounds with lower IC_50_ values were further examined for their ability to inhibit PFAR. However, because of its high IC_50_ value, the PFAR activity of Biperiden was not investigated further. Given the fact that the structural features of Biperiden are similar to those of the PFAR-active compound, Metixene, we hypothesized that Biperiden would interact with additional genes whose activity is associated with protein folding and show an inhibitory effect, which prompted us to investigate the direct interaction of BPD with *Pf*PFD (Fig. 3E).

A plausible architecture of the *Pf*PFD-Biperiden-TRiC complex was constructed using the generated structural model of the *Pf*PFD hetero-hexameric complex, and the X-Ray diffraction-based structure of the human TRiC complex (Gestaut et al., 2019) as a representative of *Pf*TRiC (Fig. 3F (i)). Biperiden was found to interact with *Pf*PFD hetero-hexameric complex via two different conformations. In conformation 1, Biperiden was found to engage at the interface of *Pf*PFD subunits 2 and 6, with free binding energy (*ΔG_bind_*) of −8.4 kCal/mol, via polar contact (H-bonds) with Glu^68^ *Pf*PFD6 with a bond length of 3.5 Å (Fig. 3F (ii)). Contrastingly, in conformation 2, Biperiden was found to interact with the core body of the *Pf*PFD complex consisting of a double beta-barrel assembly, although with a lower *ΔG_bind_* of −8.0 kCal/mol (Fig. 3F (iii)). It was, therefore, hypothesized that *Pf*PFD complexation with Biperiden in either of the conformations, would result in a diminished ability to bind and stabilize newly synthesized proteins, thereby impairing the correct folding of the nascent polypeptides. A schematic representation of complex formation between *Pf*PFD hetero-hexameric complex and Biperiden is shown.

Cellular Thermal Stability Assay (CETSA) is a method for detecting target engagement by monitoring the thermostability of a given protein in the presence of its ligand (Pantoliano et al., 2001). This is based on the principle that target-ligand interactions result in alteration in the thermodynamic parameters of the protein, affecting its stability with a corresponding increase in temperature. CETSA was optimized for binding analysis between native *Pf*PFD3 and Biperiden, which demonstrated that even at high temperatures, the treated protein fraction was more stable than the untreated fraction, indicating that Biperiden interacts with *Pf*PFD. GAPDH was used as a loading control (Fig. 3G (i) and (ii)).

To further evaluate the strength of the interaction between *Pf*PFD3 and Biperiden, we performed a competition experiment using MST, in which we used *Pf*PFD3 as the labeled target protein and, BPD, *Pf*PFD1, and *Pf*PFD2 as the unlabelled competing ligands. The concentration of *Pf*PFD3 was kept constant at 20 nM. The binding of *Pf*PFD3 to *Pf*PFD1 and *Pf*PFD2 exhibited increasing MST signals (F_norm_ [%◻] starting at 886.1 units to 892.1 units for *Pf*PFD1, and 886.4 to 893.8 for *Pf*PFD2) with increasing ligand concentration, resulting in K_D_ values of 81.6 nM and 682 nM, respectively Fig. 1C. Competing the interaction with Biperiden indicated a decrease in the MST signal with increasing ligand concentration (F_norm_ [%◻] starting at 893 unit to 888.1 units for *Pf*PFD1, and 894.5 to 887.6 for *Pf*PFD2) and deduced the K_D_ values 1.99 μM and 9.6 nM, respectively (Fig. 3H (i),(ii)). The difference in the MST signal in the presence and absence of BPD indicated that Biperiden competes for binding with the PFD subunits.

### Biperiden treatment destabilizes α-tubulin-I and PfMSP-1, substrates of the PfPFD complex

We previously reported that MSP-1 is a substrate of *Pf*PFD6 (Kumar et al., 2022a). MSP-1 is a critical molecule in the parasite egress and invasion of fresh RBCs. We evaluated the expression and localization of both α-tubulin-I and MSP-1 in the parasite to assess how Biperiden affects the function of the prefoldin subunits. Our western blot analysis revealed that the expression of both α-tubulin-I and MSP-1 gets reduced in Biperiden-treated parasites compared to the untreated ones (Fig. 4A (i), (ii), and B (i), (ii)). Confocal microscopy also showed that MSP-1 delocalizes from the surface of infected RBCs upon Biperiden treatment, as compared to control which showed surface expression of MSP-1 (Fig. 4A (iii)). NapL was taken as control (Fig. S4). These findings imply that Biperiden is a *Pf*PFD subunit inhibitor.

**Figure 4.**
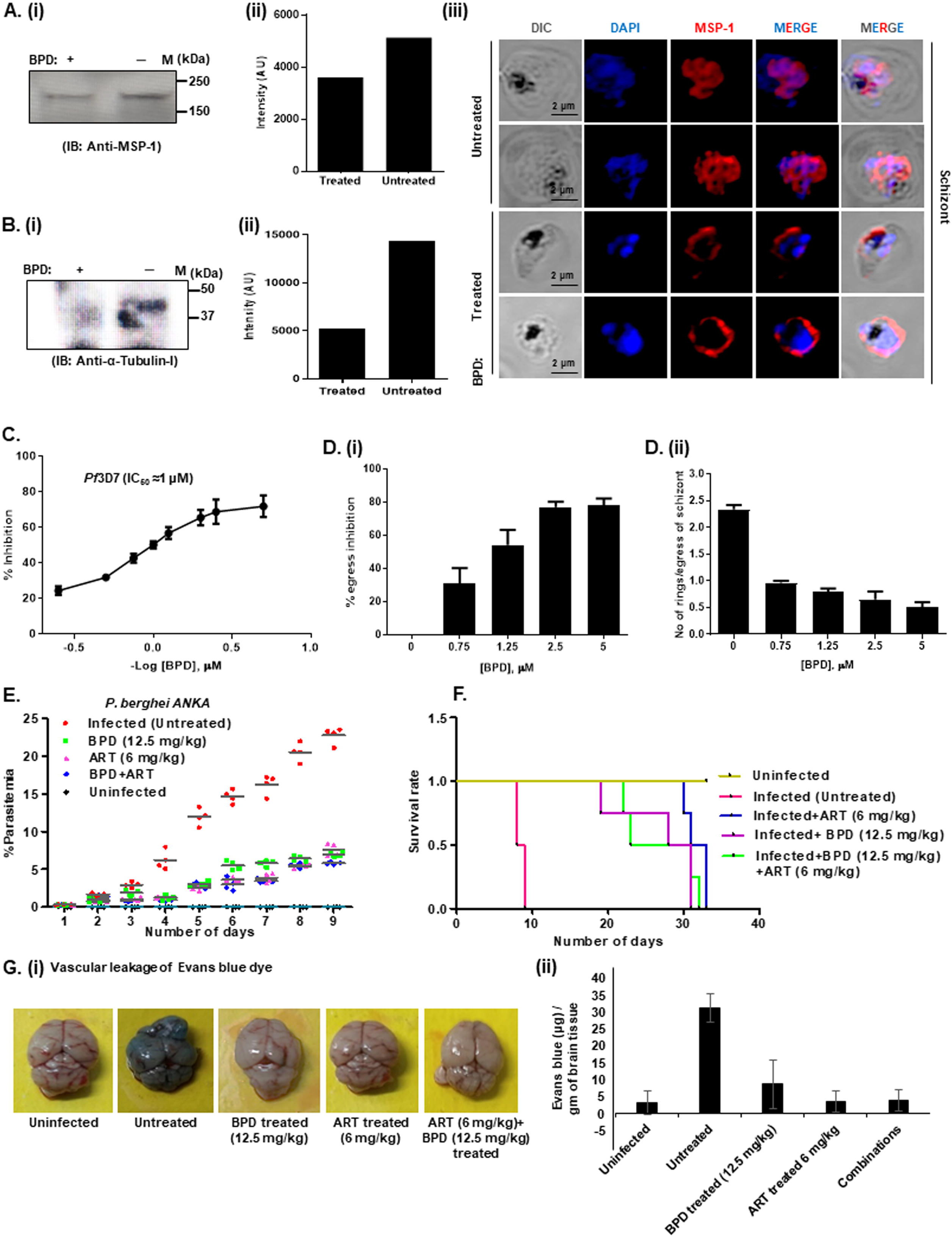
Effect of Biperiden on the PfPFD substrates and growth of malaria parasite. **(A, B) Effect of BPD on the stability of prefoldin substrates ‘MSP-1’ and α-tubulin-I**. Ring stage *Pf*3D7 parasites were treated with IC_50_ of BPD for 48 hours followed by harvesting the lysate. **(A i, ii)** Western blot analysis was conducted on total parasite lysate of treated and untreated samples using monoclonal anti-MSP-1 (1:500) and **(B i, ii)** α-tubulin-I antibodies (1:500). **(A iii)** Methanol fixed thin blood smears of treated and untreated *Pf*3D7 parasites were stained with anti-MSP-1 antisera (1:200) followed by incubation with Alexa Fluor conjugated secondary antibodies (1:200; Alexa Fluor 594, red color). DIC: differential interference contrast image, DAPI: nuclear staining using 4’,6-diamidino-2-phenylindole (blue); anti-MSP-1 antibodies (red); merge: overlay of MSP-1 with DAPI. **(C) *In vitro* anti-malarial activity of BPD**. Ring stage *Pf*3D7 was treated with varying concentrations (250 nM to 5 μM) of BPD for 72 hours. IC_50_ value was calculated by plotting the values of percent growth inhibition against log concentrations of BPD. The experiments were performed in triplicate and ± SD value was calculated for each data point. **(D) BPD treatment impaired egress and invasion of the parasite (i)** Schizont stage *Pf*3D7 was treated with varying concentrations of BPD. The relative inhibition in egress was determined by counting the number of remaining schizonts after 8 hours of treatment as compared to the untreated control (*n* = 3). **(ii)** The ability to form rings per schizont egress was determined by counting the number of schizonts and rings after 8 hours using Giemsa-stained smears (*n* = 3). **(E) *In vivo* anti-malarial activity of BPD (i)** Comparative study of percent parasitemia and their survival in *P. berghei* infected mice. Graph showing percent parasitemia in five experimental groups. On day 9, parasite load in untreated group was ~24%, while in BPD and artesunate treated group ~5% parasitemia was observed. **(ii)** Graph showing percent survival of *P. berghei* infected mice. On day 9 all mice of untreated group died while more than 50% mice survived for 20 days in the treated group. **(F) Comparison of blood–brain-barrier damage in treated and untreated mice (i)** Images of brains isolated from treated and untreated mice injected with Evans Blue dye. **(ii)** Graph showing amount of Evans Blue leakage due to damage to blood-brain-barrier in *P. berghei* infected mice.

### Biperiden treatment inhibits the *in vitro* and *in vivo* growth of malaria parasite

Next, Biperiden was investigated for its anti-malarial activity in *Pf*3D7 using an intra-erythrocytic growth inhibition assay (GIA) *in vitro*. The overall parasitemia was evaluated in comparison to the untreated control at 72 hrs. Biperiden inhibited the growth of the malaria parasite and displayed a potent anti-malarial effect, with an IC_50_ value of ~1 μM (Fig. 4C). We next evaluated the effect of Biperiden on the egress and invasion processes of the parasite. Biperiden significantly inhibited the egress of the parasite by ~60% at 1.25 μM, and ~75% at 2.5 μM and 5 μM concentrations (Fig. 4D (i)). Similarly, the number of rings formed per egress of the parasite was significantly reduced in the presence of Biperiden, as compared to control (Fig. 4D (ii)).

The anti-plasmodial activity of Biperiden was also evaluated *in vivo*, in mice infected with *P. berghei* ANKA, alone or in combination with artesunate. Group 1 included untreated infected mice, group 2 included infected mice treated with Biperiden alone (12.5 mg/kg), group 3 included artesunate-treated mice (6 mg/kg), and group 4 included mice treated with both Biperiden (12.5 mg/kg) and artesunate (6 mg/kg). On the tenth day after infection, parasitemia in untreated mice reached over 23%, and all mice died. However, it was around 7% in infected mice treated with Biperiden or artesunate alone. Parasitemia in group 4 mice treated with a combination of Biperiden and artesunate showed the highest reduction of *P. berghei* growth (~5% parasitemia) (Fig. 4E). On the twenty-third day of infection, survival of infected mice treated with Biperiden, artesunate, or a combination of Biperiden and artesunate was found to be 50% (Fig. 4F). These findings indicate that Biperiden-treated mice possess reduced parasitemia and a higher survival rate than untreated ones.

Further, an Evans Blue leakage assay was performed to determine the extent of parasite-induced blood-brain barrier disruption in infected mice. A significantly higher level of blood-brain barrier damage was observed in untreated mice compared to control (Fig. 4G (i)). Evans leakage was greater in brain tissue from untreated mice as compared to Biperiden and artesunate treated mice (Fig. 4G (ii)). These findings indicate the parasite growth-inhibitory potency of Biperiden.

## Discussion

Prefoldin complex plays an important role in maintaining protein homeostasis and cell survival. Thus alteration in its activity leads to the occurrence and development of tumors and neurodegerative diseases in humans. Thus, the abnormal expression of prefoldins is used as a biomarker to specify the prognosis of the disease (Tahmaz et al., 2022; Yesseyeva et al., 2020; Zhou et al., 2020). Owing to the significant role played by prefoldins, they are well characterized in archaea and eukaryotes. But there are no studies on *Plasmodium spp*. prefoldin proteins though they could be important for managing the parasite proteostasis. The present study characterizes the functional role of *P. falciparum* prefoldin complex in protein homeostasis and parasite survival.

Prefoldins (PFD) are the heterohexameric molecular co-chaperone, which capture nascent polypeptide (e.g. actin and tubulin) and subsequently deliver it to TRiC/CCT complex for correct folding (Gestaut et al., 2019; Siegert et al., 2000) Prefoldin is present in archaea and eukaryotes and composed of six distinct subunits (PFD1-6) that form a jellyfish-like structure. In archaea, PFD complex is comprised of two α and four β subunits, while in eukaryotes two α like (PFD3 and PFD5) and four β like (PFD1, PFD2, PFD4 and PFD6) subunits exist to form a complex (Leroux, 1999). The *Pf*PFD complex is more similar to eukaryotic PFDs since *Pf*PFD3 interacts with *Pf*PFD2 and *Pf*PFD1; and *Pf*PFD5 interacts with *Pf*PFD4 and *Pf*PFD5. These semi-complexes of *Pf*PFD3 and *Pf*PFD5 interacts with each other to form active heterohexameric complex (Liang et al., 2020).

Chaperone activity of PFD plays a significant role in restricting the protein misfolding and thus reduced toxic aggregate formation in neurodegenerative diseases. Dysfunction of PFD in humans leads to the aggregation of proteins that trigger neurodegenerative diseases like Alzheimer’s (AD), Parkinson’s (PD), and Huntington’s diseases (HD) (Sörgjerd et al., 2013; Takano et al., 2014; Tashiro et al., 2013). Similarly, *Pf*PFDs may interact with the *Pf*MSPs, specifically *Pf*MSP-1 due to the presence of intrinsically unstructured regions. We also demonstrate that *Pf*PFDs interact with *Pf*α-tubulin-I and ligulate their polymerization (Kumar et al., 2022b). Thus, chaperone activity of *Pf*PFDs is not only regulating protein homeostasis but also playing an important role in protein functioning. In line with above, in *Caenorhabditis elegans*, RNAi mediated knockdown of PFD leads to reduced α-tubulin levels; subsequently, microtubule polymerization rate is decreased (Locascio et al., 2013).

To understand the role of *Pf*PFD complex, we investigated various chemotypes from FDA and identified biperiden as a potent molecule against the complex. Biperiden inhibited formation of complex *in vitro* and *in vivo*. Moreover, biperiden also restricted interaction of *Pf*PFD complex with its substrate tubulin and led to degradation of the substrates. We further demonstrate that the biperiden has anti-malarial activity both *in vitro* and *in vivo*. Thus, Biperiden has not only helped us to understand the role of the *Pf*PFD complex but also led to development of targeted therapeutics against malaria.

Overall, this is the first study to describe the functional importance of unexplored prefoldin subunits of malaria parasite in detail and identified it as a potent pharmacological target. Our results elucidate the expression and localization of prefoldin subunits at asexual stages of malaria parasite. Supporting previously published data in archea and eukaryotes, our complex formation data further strengthen the view that prefoldin subunits exist as a complex form in *Plasmodium*. Also, we identified BPD as an inhibitor of prefoldin subunits which significantly bind to it and affect expression and localization of its substrate “MSP-1” and “α-tubulin-I” in malaria parasite. Simultaneously, our *in vitro* and *in vivo* results showed that BPD is a potent antimalarial inhibitor against malarial parasite. Collectively our results open new avenue in understating malaria biology and a step further for the development of potent antimalarial drugs.

### Limitation of the study

The study characterized the functional and biochemical roles of PfPFDs in malaria parasite. The effect of PfPFD complex attenuation on the parasite will help in identifying their substrates and detailing their role in various stages of the lifecycle. This, however, is not possible due to practical limitations since most of the PfPFDs are essential for the blood stage growth. Hence, complete genetic ablation of PfPFDs is near to impossible. Moreover, since PfPFDs function in a complex, knockout of single PfPFD will provide incomplete information about the role of this complex in parasite growth. And knockout of six genes present at different genetic loci is altogether an impossible task. Thus, the limitation of our study is partly due to the natural essentiality of these genes and also due to the practical problems associated with the current genetic modification strategies available for the malaria parasite.

## Acknowledgements

RS is UGC-SRF, Govt. of India. SG is funded by CSIR-Senior Research Associate (CSIR-SRA), VK is supported by Research Associateship Program of Department of Biotechnology, Govt. of India. AB is supported by National Post-doctoral fellowship, SERB, India (Fellowship reference no. PDF/2019/000334). This work has been funded by Drug and Pharmaceuticals Research Programe (DPRP) (Project No. P/569/2016-1/TDT) to SS. SS is a recipient of the National Women Bio scientist award from DBT.

## Author contributions

RS, VK conducted most of the experiments, analysed data and wrote the manuscript. SG: Manuscript editing and data analyses. MS: helped in conducting MST experiments JK, PM, AG, NG, HS, PV helped in biophysical experiments. RJ: did bioinformatic analysis and manuscript writing. SB: Helped in Co-immunoprecipitation assay. AB: manuscript writing. MA: Manuscript editing. SS conception of idea, data analysis, manuscript writing and approval of final draft.

## Data code availability

The article includes all datasets generated or analyzed during this study.

Data reported in this paper will be shared by the lead contact upon reasonable request.

## STAR methods

### In vitro culture of *P. falciparum*

*Plasmodium falciparum* 3D7 parasite culture was grown in complete RPMI 1640 media supplemented with AlbumaxII (Gibco, USA), hypoxanthine (Sigma-Aldrich, MA, USA) and gentamycin (Gibco, USA) using O^+^ human erythrocytes. Culture was maintained at ~5% parasitemia and 2% haematocrit in 37°C incubator in a mixed gas (5% CO_2_, 5% O_2_ and 90% N2) chamber. Synchronization of parasites was done by treatment of culture with 5% sorbitol followed by 65% percoll. Cell pellets were extensively washed twice with incomplete RPMI media before placing into culture. Mixed stage and synchronized cultures were harvested and parasite pellet stored at −80 °C for future experiments.

### Cloning, expression and purification of PfPFDs of *Pf*3D7

Genes for all six prefoldin subunits of *Pf*3D7 [PF3D7_1107500 (*Pf*PFD), PF3D7_1416900 (*Pf*PFD2), PF3D7_0718500(*Pf*PFD3), PF3D7_0904500 (*Pf*PFD4), PF3D7_1128100 (*Pf*PFD5) and PF3D7_0512000 (*Pf*PFD6)] were cloned in pET28a vector (Novagen, Merck KGaA Madison, WI, USA) using BamHI and XhoI restriction sites and expressed in *E. coli* BL21 (DE3) cells. Prefoldin subunits were PCR amplified from cDNA of *Pf*3D7 using gene specific primers. Ni-NTA (Qiagen, Hilden, Germany) affinity purification of all Prefoldin recombinant protein was carried out in lysis buffer (50 mM Tris/HCl, 300 mM NaCl and 0.02% Na-azide, pH 8.0).

### Generation of antisera against PfPFDs

Polyclonal antibodies were raised against all six prefoldin subunits in Balb/C mice following standard protocol by using purified recombinant proteins as an immunogen. Briefly, mice were immunized with emulsion containing 1:1 ratio (v/v) of immunogens (50 μg) and freund’s complete adjuvant. Two booster doses were administered with 1:1 ratio (v/v) of immunogens (25 μg) and freund’s incomplete adjuvants before collecting mice sera. Antibodies were checked for specificity by performing western blotting before performing experiments.

### Generation 3D-structure model of *Pf*PFD complex

Amino acid sequences of the PFD subunits 1-6 from *P. falciparum* strain 3D7 (1: PF3D7_1107500; 2: PF3D7_1416900, 3: PF3D7_0718500, 4: PF3D7_0904500, 5: PF3D7_1128100, and 6: PF3D7_0512000) were retrieved from the PlasmoDB database (https://plasmodb.org/plasmo/app) (Aurrecoechea et al., 2009). A multiple threading approach, which is one of the most common structure prediction methods in structural genomics and proteomics, was employed to generate 3D-structural coordinates of the *Pf*PFD subunits. To accomplish this feat, individual structural models of each subunit were generated using I-TASSER (Iterative Threading ASSEmbly Refinement), a web server that uses a hierarchical approach to protein structure prediction and structure-based function annotation (https://zhanggroup.org/I-TASSER/) (Zhang, 2008), as described previously (Yadav et al., 2019). Structural models of *Pf*PFD subunits 1-6, thus generated, with higher values of Confidence-score (C-score) were selected, and subjected to structural refinement by using ModRefiner (https://zhanglab.ccmb.med.umich.edu/ModRefiner/) which is an algorithm-based approach for atomic-level, high-resolution protein structure refinement (Xu and Zhang, 2011). The refined structural models of *Pf*PFD subunits were rendered with PyMOL Molecular Graphics System, v2.1 by Schrödinger, LLC (http://pymol.org/2/) (Delano, 2002), and set to submit to generate *Pf*PFD hetero-hexameric complex structure.

The X-Ray diffraction-based structural model of the human TRiC (T-complex protein Ring Complex, also known as Chaperonin Containing TCP-1 (CCT))-PFD complex (PDB ID: 6NR8; resolution: 7.80 Å) (Gestaut et al., 2019) was used as a suitable template to generate a 3D structural model of the *Pf*PFD hetero-hexameric complex, as described previously (Jain et al., 2020a). The reliability of the *Pf*PFD structural model was assessed by examining backbone dihedral (torsion) angles: phi (Ø) and psi (Ψ) of the amino acid residues lying in the energetically favorable regions of Ramachandran space (Ramachandran and Sasisekharan, 1968). This was done by using PROCHECK v.3.5 which checks the stereochemical quality of a protein structure, producing a number of PostScript plots analyzing its overall and residue-by-residue geometry (https://www.ebi.ac.uk/thornton-srv/software/PROCHECK/) (Laskowski et al., 1993). Percent quality measurement of the protein structures was evaluated by using four sorts of occupancies called ‘core’, ‘additional allowed’, ‘generously allowed’, and ‘disallowed’ regions. The 3D structural model of *Pf*PFD, thus generated, was subsequently used for *in silico* and *in vitro* interaction analysis, and inhibition studies.

### Co-Immunoprecipitation Assay for interaction PfPFD with other subunits and its substrates

Co-immunoprecipitation assay was performed using Pierce Co-immunoprecipitation (Co-IP) Kit to confirm the existence of PFD as a complex form in malaria parasite. Briefly, 70 μg anti-PFD6 antibodies were cross linked to AminoLink plus coupling beads followed by extensive washing with wash buffer. Mixed stage culture (~8% parasetemia) was subjected to saponin lysis (0.15%) followed by RIPA lysis. Parasite lysates were pre cleared with control beads before using it. Beads crosslinked with antibodies were incubated with RIPA lysed mixed stage parasite lysate overnight at 4°C. Sample was eluted by using elution buffer and divided into six groups. Eluted samples along with appropriate control were resolved on 12% SDS-PAGE and transferred to nitrocellulose membrane. Blots were probed with anti-prefoldin subunits antisera separately in 1:1000 dilution (*Pf*PFD1, *Pf*PFD2, *Pf*PFD3, *Pf*PFD4, *Pf*PFD5 and *Pf*PFD6). HRP conjugated anti-mice antibodies (1:5000) were used as a secondary antibodies (Sigma Aldrich,USA). Blots were developed using diamino benzidine/H_2_O_2_ substrate (Sigma-Aldrich, MA, USA).

A similar protocol was followed to investigate the interaction of *Pf*PFD2 with α-tubulin-I. Anti-PFD-2 sera were crosslinked to the AminoLink plus coupling beads. Anti-α-tubulin-I (1:1000) antibodies were used for western blotting. To further validate this interaction, reverse Co-IP was performed, where anti-α-tubulin-I antibodies were cross linked and anti-PFD2 antibody were used for western blotting.

Immunoprecipitation was performed where *Pf*PFD2 and *Pf*PFD6 antisera was crosslinked to the AminoLink plus coupling beads beads. Eluted sample was probed with primary antibodies MSP-I (1:2000) and α-tubulin-I (1:800) respectively followed by probing with secondary anti-rabbit and anti-mice (1:5000) antibodies respectively..

### MicroScale Thermophoresis assays

The kinetic measurements of one to one binding of prefoldin subunits were conducted in NanoTemper Monolith NT.115 instrument. Briefly, 20 μM of *Pf*PFD3 and *Pf*PFD5 were labeled using NanoTemper’s Protein Labelling Kit RED-NHS (L001, NanoTemper technologies, Germany) (Akshay Munjal, Deepika Kannan, 2022). The concentration of labelled prefoldin subunit proteins were kept constant at 20 nM. The unlabeled *Pf*PFD1 and *Pf*PFD2 for *Pf*PFD3, *Pf*PFD4, *Pf*PFD5 and *Pf*PFD6 in varying concentrations (*Pf*PFD1-7.5 uM, *Pf*PFD2-28 uM, *Pf*PFD3-20 nM, *Pf*PFD4-20 uM, *Pf*PFD5-13.4 uM, *Pf*PFD6-20 uM) were serially titrated in decreasing concentration in 1XPBS (pH 7.5) with 0.01% tween-20 in 1:1 dilution. For measurements, samples were filled into the capillaries (K002 Monolith NT.115) and thermophoretic mobility was analyzed.

Competition assay was performed by NanoTemper Monolith NT.115 instrument to check whether the binding of a drug hinders the interaction between those PFD subunits. For this experiment, stock solution containing labelled *Pf*PFD3 concentration 20 nM was mixed with *Pf*PFD1(7.5 uM), and *Pf*PFD2(28 uM) incubated for 10-15 minutes. At this concentration protein is in their maximum bound state. *Pf*PFD3-*Pf*PFD1, *Pf*PFD3-*Pf*PFD2 complex was mixed with a serial dilution of drug BPD in PBS buffer with 0.01% tween-20 at starting concentration of 100 μM and measured under the same condition as in the absence of drug experiments.

### Real time PCR analysis of PfPFDs

Expression of prefoldin subunits was evaluated at transcript level in asexual blood stages of *Pf*3D7 using real time PCR (StepOnePlus Real time PCR system Applied Biosystems, USA). The 18S rRNA gene of *Plasmodium falciparum* was chosen as a positive control. Reaction mixture (10-μl) comprises 1 μl cDNA (1 μg), 5 μl SYBR™ Green PCR Master Mix (Applied Biosystems™), and 1 μl (5 mM) prefoldin subunits specific forward and reverse primers. The PCR conditions include initial denaturation at 95°C for 5 min, followed by amplification for 40 cycles of 15 seconds at 95°C, 5 seconds at 55°C, and 1 minute at 72°C, with fluorescence acquisition at the end of each extension step. Amplification was immediately followed by a melt program consisting of 15 seconds at 95°C, 1 min at 60°C, and a stepwise temperature increase of 0.3°C/s until 95°C, with fluorescence acquisition at each temperature transition.

### Immuno-Fluorescence Assay (IFA)

Thin smear of mixed stage parasite culture was fixed in ice cold methanol for 30 minutes at −20°C. Fixed smear was permeabilized with PBS/Tween-20, and blocked in 5% of BSA (w/v) in 1xPBS with for 2 hours at room temperature. For localization studies, primary antibodies of prefoldin subunits (1:200) were added followed by incubation at room temperature for 2 hours. Alexa Fluor 488 conjugated anti-mice (1: 250, green colour, Molecular Probes, Invitrogen, Carlsbad, CA, USA) was used as secondary antibody. For colocalization studies primary antibodies i.e, anti-prefoldin subunits antibodies (1:200), anti-rabbit *Pf*NapL (1:250) anti-rabbit MSP1 (1:250), anti-rabbit α-tubulin-I (1:250) were used. Alexa Fluor 488 conjugated anti-mice (1: 250, green colour, Molecular Probes, Invitrogen, Carlsbad, CA, USA) and Alexa Fluor 546 conjugated anti-rabbit (1:250, red colour; Molecular Probes) were used as secondary antibodies. DAPI-antifade (Invitrogen, Life technologies corporation, Eugene, OR, USA) was used to counterstain parasite nuclei followed by mounting of slides with cover slips. The slides were viewed under confocal microscope at 100X magnification (Olympus Corporation, Tokyo, Japan).

### Stage specific expression of PfPFDs by Immunoblot analysis

Mixed stage asexual cultures of *Pf*3D7 (parasitemia 5–10%) were subjected to saponin lysis (0.15% w/v) followed by extensive washing with 1x PBS. Lysed parasite pellets (10 μg total protein), recombinant purified proteins (positive control), infected RBC cytosol, uninfected RBC pellet, and crude extract of *E. coli* (negative control) were resolved on 12% SDS/PAGE and transferred to NC membrane. The transferred blots were blocked in 5% BSA in 1x PBS overnight at 4 °C, and probed with mice anti-prefoldin subunits antisera (1 : 5000) followed by incubation with horseradish peroxidase (HRP) –conjugated goat anti-mice IgG (1: 2000). Blots were developed using diamino benzidine/H_2_O_2_ substrate.

To check stage specific expression of prefoldin subunits, western blotting was performed at ring, trophozoite and schizont stages of malaria parasite. Synchronized ring, trophozoite, and schizont stage cultures of *Pf*3D7 were harvested separately and subjected to saponin lysis. Parasite pellets were incubated with RIPA lysis buffer for 30 min at 4°C to break the parasites membrane and release its cytosolic content. Supernatant of lysed parasite of all three stages (10 μg total protein) was resolved on 12% SDS/PAGE and transferred to NC membrane. The blot was blocked in 5% BSA in PBS overnight at 4 °C, and probed with mice prefoldin subunits antisera (1:5000) followed by incubation with horseradish peroxidase (HRP) –conjugated goat anti-rabbit IgG (1:2000). Blots were developed using diamino benzidine/H_2_O_2_ substrate.

### Enzyme Linked Immune Sorbent Assay (ELISA) to study interaction of PfPFDs with α-tubulin-I

ELISA 96 plate wells were coated with 100 nanograms of purified *Pf*PFD1, *Pf*PFD2, *Pf*PFD3, *Pf*PFD4, *Pf*PFD5 and *Pf*PFD6 bait proteins and blocked overnight at 4°C with 5% BSA in PBS, followed by incubation of plate at room temperature for 2 hours with increasing titre ranges from 0-500 ng of prey protein α-tubulin-I. After washing with 1XPBS primary prefoldin antisera and secondary anti-mice HRP conjugated antibody were incubated at room temperature for 2 hours. After washing with 1XPBS detection reagent TMB (HIMEDIA) was added, after development of colour stop solution (3M HCl) was added to the reaction. Absorbance was measured in microplate reader at 450 nm. Graph was made using Graphpad PRISM software.

### Surface Plasmon Resonance (SPR)

Interaction of *Pf*PFD2, *Pf*PFD1 and *Pf*PFD3 with α-tubulin-I of *Plasmodium falciparum* were evaluated by Auto LAB ESPRIT SPR instrument (Kinetic Evaluation Instruments BV, The Netherlands). Briefly, 25 μM purified recombinant *Pf*PFD2 proteins were immobilized on gold sensor chip that had been activated through amine coupling. 1x PBS was used as immobilization and binding buffer. α-tubulin-I was injected in increasing concentration (100 nM, 250 nM, 500 nM, 750 nM, 1 μM) over immobilized chip surface. 50 mM NaOH buffer was used to renew the chip surface and Auto Lab ESPRIT kinetic evaluation software was used to assess the data. Similar protocol was followed for negative control experiment where *Pf*PFD1 and *Pf*PFD3 was immobilized.

### LOPAC^®1280^ library screening for a possible inhibitor of *Pf*PFD protein folding activity

In a recent investigation by Aline Bamia *et al*., novel small molecules with Prion (PrP^Sc^) propagation-inhibitory activities were identified, which interfered with the Protein Folding Activity of the Ribosome (PFAR), and significantly prolonged the survival of prion-infected mice (Bamia et al., 2021). Using an *in silico* therapeutic repositioning approach, we screened LOPAC^®1280^ Library of 1,280 Pharmacologically Active Compounds (Sigma-Aldrich) based on similarities with one of the potent PFAR inhibitors identified in the study, Metixene (https://www.sigmaaldrich.com/IN/en/product/sigma/lo1280). Metixene is a member of piperidines and has a role as an antiparkinson drug and a muscarinic antagonist. Structural Data Format (SDF) files of Metixene and LOPAC^®1280^ library were retrieved from PubChem, a database of freely accessible chemical information of chemical molecules and their activities against biological assays (https://pubchem.ncbi.nlm.nih.gov/), and Sigma-Aldrich, respectively. Structural superimposition of LOPAC^®1280^ ligands with Metixene was done by using Discovery Studio Visualizer v20.1.0.19295, developed by Dassault Systèms Biovia Corp. (https://www.3ds.com/products-services/biovia/), and overlay structural similarity for each of the 1,280 compounds were evaluated, taking Metixene as a reference compound. Structural superimposition analysis was done by using ChemDraw Ultra v12.0.2.1076, one of the CambridgeSoft products for producing a nearly unlimited variety of biological and chemical drawings (https://perkinelmerinformatics.com/products/research/chemdraw).

### *In silico* interaction analysis of Biperiden with *Pf*PFD

Structural Data Format (SDF) file of Biperiden.HCl was retrieved from PubChem, converted to standard PDB format, followed by the generation of its energy-minimized 3D-structural model by using Chem3D Pro 12.0, as described previously (Dangi et al., 2019; Yadav et al., 2019). Molecular docking studies were performed by using Autodock Vina Tools 1.5.6 to rationalize the inhibitory activity of Biperiden against *Pf*PFD (Dangi et al., 2019; Jain et al., 2020b; Srivastava et al., 2019; Trott oleg and Arthur J. Olson, 2010; Yadav et al., 2019). We ensured that the entire *Pf*PFD complex was covered while constructing a virtual 3D grid for the *in silico* interaction analysis. A grid of 80 × 100 × 80 with x, y, and z coordinates of the center of energy, 209.514, 147.278, and 209.119, respectively was constructed through the Autogrid module of AutoDock Tools, with default spacing. Top scoring docked conformations of Biperiden were selected based on their most negative free binding energies and visualized for polar contacts (H-bonds; if any) with the amino acid residues of *Pf*PFD complex using PyMOL Molecular Graphics System (Delano, 2002).

### CETSA (Cellular Thermal Shift Assay)

To confirm the binding of drug molecule BPD with *Pf*PFD3 cellular thermal shift assay (CETSA) was performed, a process to assess the binding of ligand molecule with the target protein in cell lysate. To perform this, synchronized late trophozoite stage 8% parasitized RBCs were treated with 5 μM concentration of BPD. After six hours of treatment cells were pelleted down and lysed with 0.05% saponin followed by washing with ice-cold 1X PBS and lysis with RIPA. Lysate was cleared by centrifugation at 13000 rpm for 30 minutes at 4°C. After that, soluble fraction of parasite lysate again treated with BPD and kept at room temperature for 30 minutes. sample was heated at varying temperature from 4°C-60°C for 3-5 minutes followed by incubation at RT for 5 minutes. After centrifugation, sample was resolved on SDS-PAGE and transferred to nitrocellulose membrane. Transferred blot was probed with primary *Pf*PFD3 antibody (1:500) and secondary anti-mice antibody (1:5000).

### Effect of BPD treatment on substrates of PfPFDs within parasite

Substrate inhibition assays were performed to check the effect of BPD on interacting partners. Briefly, ring stage parasites (0.8% parasitemia, 3% haematocrit) were treated with BPD at a concentration equivalent to its IC_50_ (1 uM) for 48 hours. Thin blood smears of treated culture were prepared on glass slides for IFA experiments while, rest of parasite culture was harvested for western blotting. IFA and western blotting were performed according to above described protocol.

### Parasite Growth inhibition assays

Parasite growth inhibition assays were performed to evaluate the effect of Biperidin hydrochloride on parasite. Briefly, synchronized culture of *Pf*3D7 (90 μl) with over 80% ring stage (0.8 % parasitemia, 2% haematocrit) was treated with drug (10 μl) in increasing concentration (250 nm, 500 nM, 750 nM, 1 μM 2.5 μM and 5 μM). Dilution of Biperiden hydrochloride was prepared in incomplete RPMI. The microplates were maintained at 37°C in a controlled atmosphere (5% O_2_, 5% CO_2_ and 90% N_2_) for 72 hours. Giemsa-stained thin blood smears of *P*. *falciparum* were made and ~3000 red blood cells (RBCs) counted in light microscopy. Graph was made using Graphpad PRISM software. Experiment was performed in triplicates. Percent growth inhibition was calculated by using the formula:

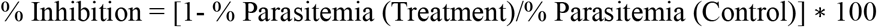

### Parasite Egress and Invasion Assay

In order to determine the effect of BPD on invasion and egress rate of parasite, late schizont (~45-47 h post invasion (hpi)) were diluted to ~4% of parasitemia and 2% of hematocrit. Parasite was treated with varying concentrations of BPD (5 μM, 2.5 μM, 1.25 μM, 0.625 μM). Untreated parasite was taken as control. After 8 hours of treatment thin smear was made on glass slides and stained with Giemsa. Approximately 3000 RBCs were counted under light microscope at 100X magnification. Percent egress was calculated by the formula: (number of schizont before treatment− No of schizont in treated sample)/(number of schizont before treatment-number of schizont in untreated sample treatment) *100. Initial number of schizont were taken as 100%. Percent egress inhibition was calculated as:100-percent egress. Number of rings formed per schizont egress = (Number of rings)/(Number of schizonts before treatment-number of schizonts after treatment) (Garg et al., 2013; Raj Kumar Sah, Swati Garg, Poonam Dangi, Kalaiarasan Ponnusamy, 2019).

### Infection of mouse with *P. berghei* ANKA and monitoring of parasites

BALB/c female mice of six weeks age were divided into four groups and each group consisted of four animals. At day 0, mice were infected intra peritoneally with 1×10^6^ infected red blood cells (100 μl, diluted in PBS), which was obtained from *P. berghei* ANKA infected donor mice. BPD and artesunate were dissolved in a solvent mixture of DMSO and PBS. Group 1 mice were treated with artesunate at a concentration of 6 mg/kg (positive control), Group 2 was treated with 12.5 mg/kg of BPD, while group 3 was treated in combination of 12.5 mg/kg BPD and 6 mg/kg artesunate. Group 4 was left as untreated (negative control). Blood samples from tail end of infected mice were taken on a daily basis for determination of parasitemia. Parasitemia was determined by observing the giemsa stained smears under microscope at 100× magnification (Olympus Corporation, Tokyo, Japan).

### Evans Blue dye leakage assay

Evans Blue solution (2%) prepared in PBS was injected intraperitoneally (4 ml/kg of mice) in infected and uninfected mice. After 24 h of stain circulation, brain from one mouse from each group were removed under anaesthetized condition (by ketamine-xylazine cocktail, Sigma-Aldrich). Mice brains were placed in dimethyl formamide (Sigma-Aldrich) followed by incubation at 56°C overnight. Absorbance of the supernatant was measured at 610 nm in a microplate reader (Varioskan Flash, Thermo Fisher Scientific, USA). The dye leakage was calculated through extrapolation from the standard curve prepared for varying concentrations of evans blue.

### Animal handling

Animal studies were performed in accordance with guidelines of the Institutional Animal Ethics Committee (IEAC) of Jawaharlal Nehru University (JNU), Delhi and Committee for Control and Supervision of Experiments on Animals (CPCSEA).

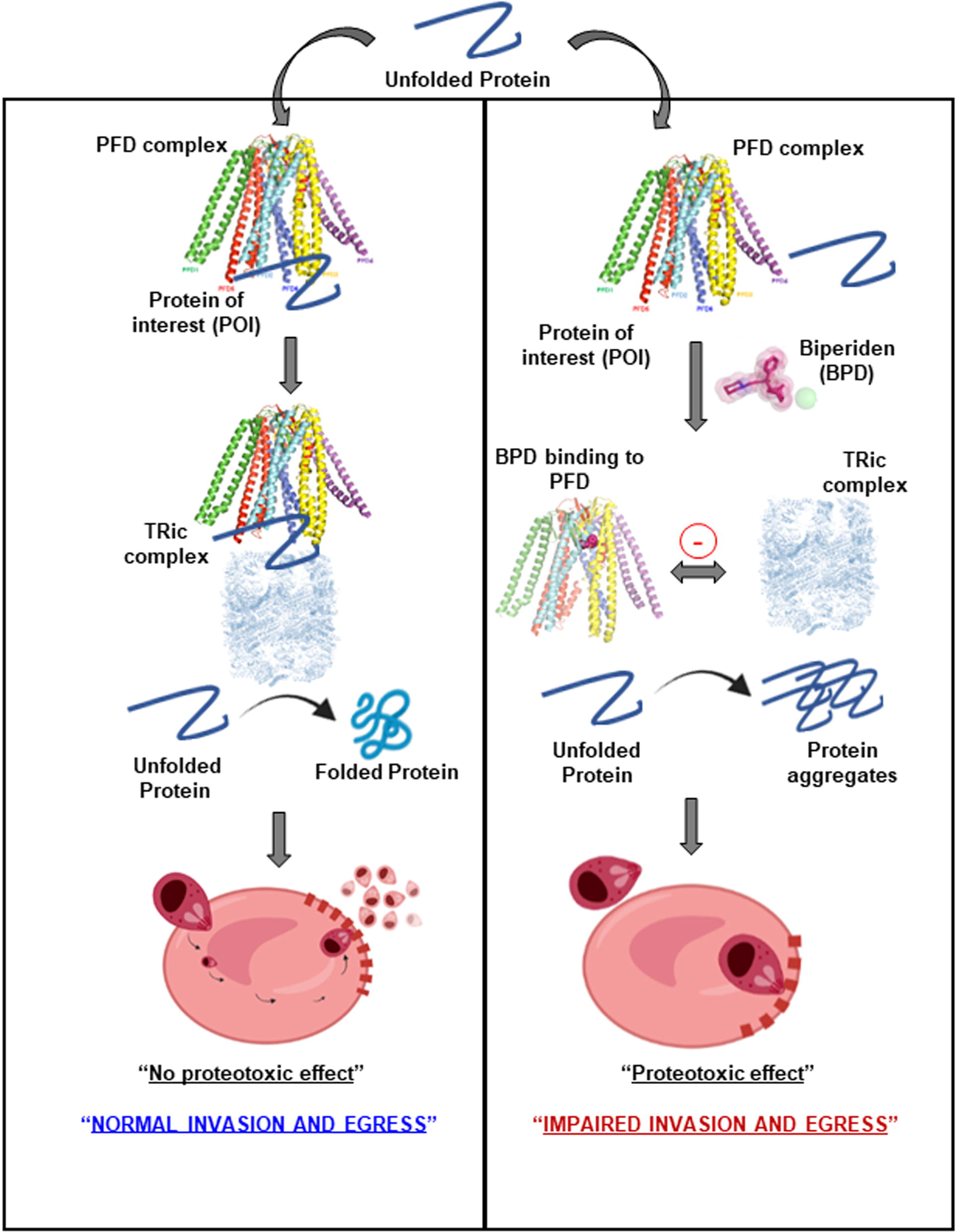

