## Supplementary file for "Chaperonin activity of *Plasmodium* prefoldin complex is essential to guard proteotoxic stress response and presents a new target for drug discovery"

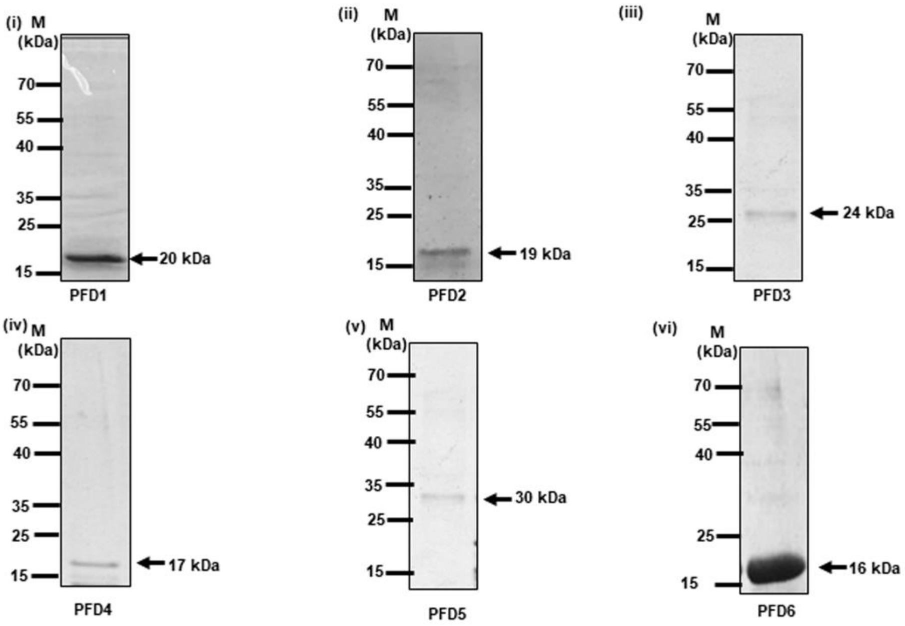


**Fig. S1: Expression and purification of prefoldin subunits of *Pf*3D7.** SDS-PAGE of purified recombinant prefoldin subunits. *Pf*PFD1 (i), *Pf*PFD2 (ii), *Pf*PFD3 (iii), *Pf*PFD4 (iv), PfPFD5 (v), PfPFD6 (vi).

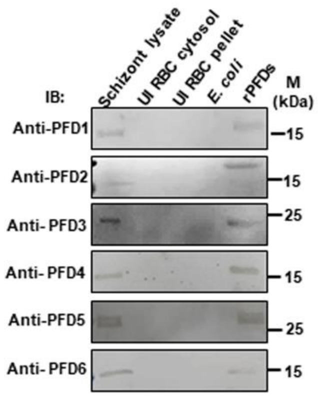


**Fig. S2: Evaluation of prefoldin subunits antibody specificity.** Western blot analysis of *Pf*PFD1, *Pf*PFD2, *Pf*PFD3, *Pf*PFD4, *Pf*PFD5 and *Pf*PFD 6 in total parasite lysate using their specific antibodies.

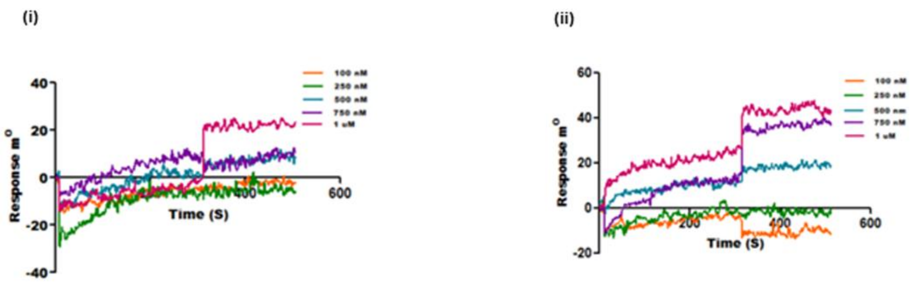


**Fig.S3 Interaction of prefoldin subunits with α-tubulin-I.** Purified recombinant *Pf*PFD1 and *Pf*PFD3 subunits (25 μM) were immobilized on gold sensor chip followed by titration with varying concentrations of α-tubulin-I. The net dissociation constant graph shows no affinity (ii and iii).

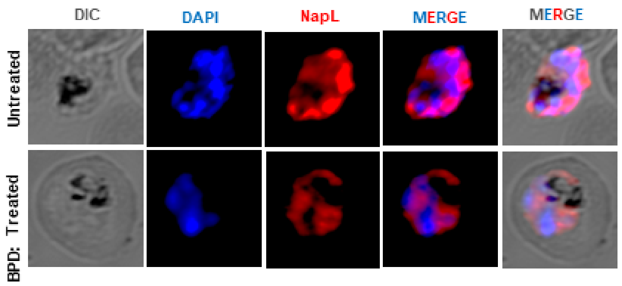


**Fig.S4 Effect of BPD on cellular localization of NapL.** Methanol fixed thin blood smears of treated and untreated *Pf*3D7 infected erythrocytes were stained with anti-NapL antisera (1:200) followed by incubation with Alexa Fluor conjugated secondary antibody (1: 200; Alexa Fluor 594, red color). DIC: differential interference contrast image, DAPI: nuclear staining using 4',6-diamidino-2-phenylindole (blue); anti-NapL antibodies (red); merge: overlay of NapL with DAPI.
